# Exploring influenza A virus receptor distribution in the lactating mammary gland of domesticated livestock and in human breast tissue

**DOI:** 10.1101/2025.04.16.649193

**Authors:** Rahul K. Nelli, Tyler A. Harm, Bailey Arruda, Chris Siepker, Olufemi Fasina, Jennifer M. Groeltz-Thrush, Amy Baker, Rachel Phillips, Brianna Jones, Virginia Espina, Hannah Seger, Paul J. Plummer, Todd M. Bell

## Abstract

The spread of the highly pathogenic avian influenza (HPAI) H5N1 virus among dairy cattle illustrates the adaptability of influenza A viruses (IAV) to infect non-traditional species. While IAV-specific sialic acid (SA) receptors have been identified in the mammary glands of dairy cattle, their presence in pigs, sheep, goats, and alpacas has not been studied until now. The zoonotic transmission of HPAI H5N1 to dairy and poultry farm workers during outbreaks raises public health concerns. This study employed lectin histochemistry to examine the mammary glands of livestock and humans. We found that these tissues were rich in SA α2,6-Gal receptors, followed by SA α2,3-Gal receptors, essential for IAV binding. Notably, the A(H5N1) clade 2.3.4.4b virus could bind to mammary tissue from both cattle and pigs. These findings highlight the potential for HPAI H5N1 to infect and spread within the mammary glands of production animals and humans.

## INTRODUCTION

Since the first human case of highly pathogenic avian Influenza (HPAI) H5N1 was reported in 1997, there has been limited evidence to suggest human-to-human transmission. With spillover from birds to humans, there have been years in which the peak annual human cases worldwide reached <150 ^1–4^. The case fatality ratios were close to 50%, raising serious public health concerns, albeit with low overall reported case numbers ^2,4,5^. Interestingly, the current outbreak of HPAI H5N1 in dairy cattle and poultry in the U.S. has resulted in numerous spillovers into farm workers handling HPAI H5N1-affected dairy herds and poultry flocks. As of March 7, 2025, there have been 70 CDC-confirmed cases, with one reported death associated with HPAI H5N1 ^6,7^. Influenza A viruses are known to replicate well in the respiratory and intestinal tract of animals and humans ^8,9^. However, the continued HPAI H5N1 viral replication in the mammary gland of novel species like dairy cattle ^10,11^ with a combination of continued spillover into numerous naive mammalian species ^12^, raises the concern of HPAI H5N1 virus adaption to mammalian species with a possibility of continued mammal-to-mammal and even human-to-human spread.

To assess the current feasibility of HPAI H5N1 viral replication within the mammary glands of other exposed animals, this study aimed to delineate the spatial distribution of influenza A virus (IAV) specific sialic acids (SA) receptors in the mammary glands of various production animals: pigs, sheep, goats, cattle, and alpacas in comparison to human breast tissue. Due to recent spillovers from production animals into people, this study also examined human breast tissue.

The mammary gland is a remarkable structure responsible for producing milk, a nutrient-rich fluid that nourishes. The importance of the mammary gland cannot be overstated, as it ensures species continuity and offspring health and development. It also contains antibodies that help to protect the young from infections ^13,14^. Most humans depend on milk, or its byproducts, for this highly nutritious fluid that contains essential proteins, carbohydrates, fats, vitamins, and minerals ^15^. The mammary gland’s structure is complex and consists of a network of alveolar lobules interconnected by ducts and supported by connective, adipose, and smooth muscle tissues (Figure 1). The lobules are responsible for milk production, while the ducts transport the milk to the cisterns for suckling. The mammary gland undergoes significant hormonal and metabolic changes during pregnancy and lactation, adapting to the demands of milk production and the growth of the fetus ^16^. The milk composition can vary among different species, reflecting the specific needs of their offspring ^17^.

**Figure 1:**
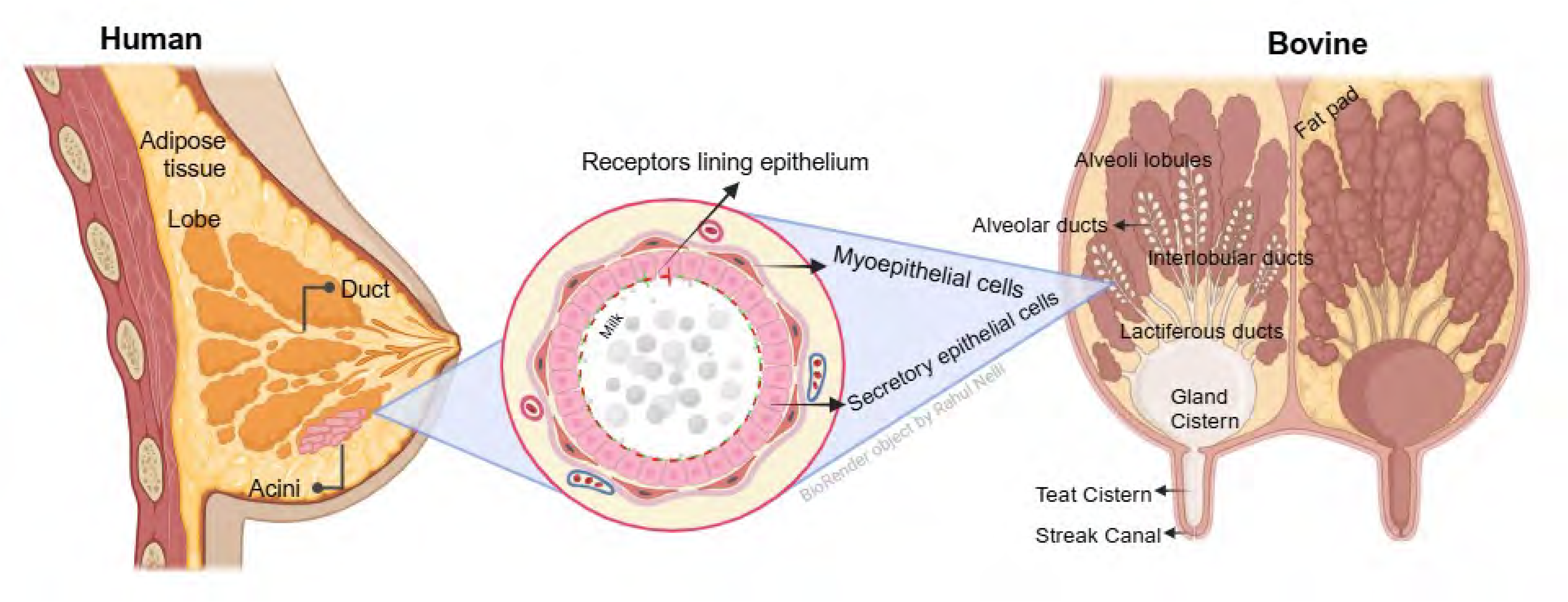
Illustration of sagittal section anatomy of the mammary gland of a cow. Inset image showing the microscopic anatomy of alveolar lobules with secretory epithelial cells, myoepithelial cells, and milk secretions.

However, the mammary gland is not immune to infections. Pathogens, such as bacteria, viruses, and fungi, can invade the mammary gland and cause mastitis, a painful and debilitating condition. Interestingly, retrograde infection is the most common route or method of infection, where microorganisms enter through the teat opening and into the canal and cistern ^18^. In dairy cows, mastitis in animals can lead to decreased milk production, reduced quality, and even death in severe cases ^19^. The infection can spread to other animals through contaminated equipment or by direct contact, posing a significant threat to the health of an entire herd ^20^. The consequences of mastitis are far-reaching. In addition to causing pain and suffering for the affected animal, mastitis can have a negative impact on the economic viability of livestock production. This includes reduced milk production, increased veterinary costs, and the potential for culling infected animals, which can result in significant financial losses for farmers. Moreover, mastitis can lead to transmitting pathogens to humans through contaminated unpasteurized milk, posing a public health risk.

The recent discovery of HPAI H5N1 clade 2.3.4.4b virus in the mammary gland and milk of dairy cattle sets a new scientific precedent, emphasizing the adaptability of H5N1 HPAI viruses to non-avian species and its potential for cross-species transmission, including to humans. A key factor in the susceptibility to HPAI H5N1 infection is the presence of specific host cell receptors that the virus uses for entry ^21^. Influenza A viruses utilize host SA for initial attachment and entry into cells ^22^. Sialic acids are also potential receptors or co-receptors for other viral families such as *Coronaviridae* ^23^, *Paramyxoviridae*, *Flaviviridae* ^24^, *Picornaviridae* ^25^, *Reoviridae* ^26^, *Parvoviridae* ^27^, *Adenoviridae* ^28^, *Papillomaviridae* ^29^, *Polyomaviridae* ^30^, and *Caliciviridae* ^31^. In addition, many bacteria, fungi, and parasites exploit sialic acids as growth substrates for immune evasion, host cell entry, and enhanced virulence, including those that cause mastitis, such as *E. coli*, *Streptococcus sps*., *Pasteurella sps*., *Candida sps.*, *Aspergillus fumigatus*, *Leishmania* and *Trypanosomes* ^32–36^.

Also sialic acids are also one of the major components of milk and play an important role in human nutrition, particularly in brain development ^37,38^. Cattle and goat milk predominantly contain sialic acid forms N-glycolylneuraminic acid (Neu5Gc) versus N-acetylneuraminic acid (Neu5Ac), and their content decreases with the progression of the lactation stage ^39,40^. Also, sialic acid levels vary depending on the diet of the animal and have been shown to regulate the microbiota-gut-mammary axis during mastitis of Holstein cows ^41^. In addition, Neu5Ac was also one of the key metabolites that was found to be elevated in metastatic breast tumors in humans ^42^. Thus, suggesting that SA levels may vary depending on the lactational stage, diet, and underlying disease conditions.

Therefore, understanding the sialic acid interactions in a complex organ like a mammary gland with frequent interactions with microbial and hormonal changes during the lactational stage is important.

## MATERIALS AND METHODS

### Sample collection

This study was approved by the Institutional Animal Care and Use Committee (IACUC) of the College of Veterinary Medicine, Iowa State University, Ames, Iowa (IACUC-24-090). In the case of the human mammary gland, consultation and approvals from ISU and GMU [Institutional Review Board approval (GMU IRB# 478170 and 477703)] were obtained before using the archived tissue blocks. Three animals of each species studied were procured from the Iowa State University Research farms (sheep, goats, beef cattle) from approved vendors (alpacas), while archived tissues for the rest (Table 1). Following euthanasia, formalin-fixed and paraffin-embedded sections of the mammary gland from the area of the teat cistern, gland cistern, interlobular (collecting) duct, and secretory alveoli (glandular epithelium) were collected from these animals (Figure 2).

**Figure 2:**
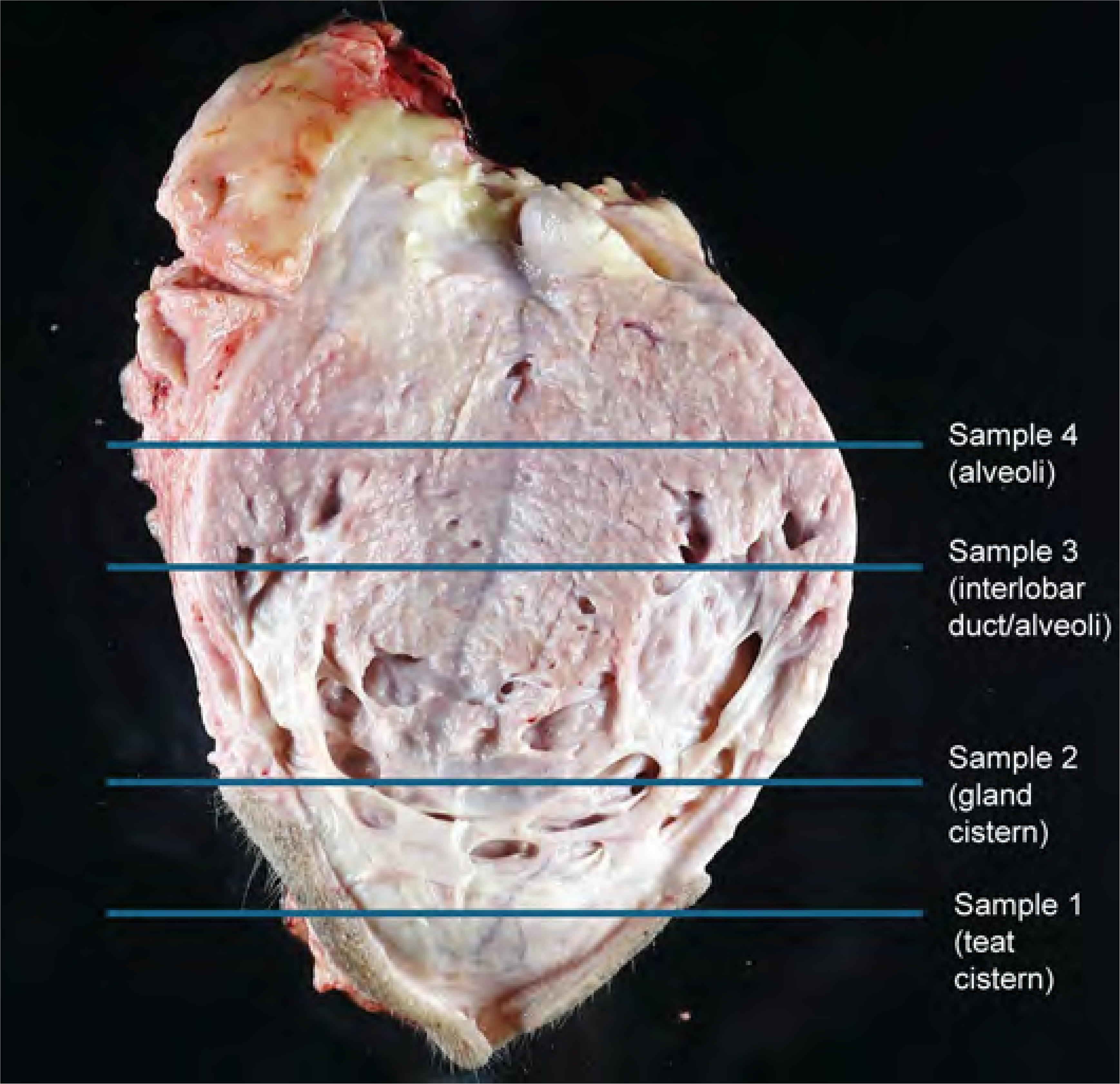
Cross-sectional gross anatomy of the mammary gland of a cow. Lines indicate representative areas where samples were collected for histological studies.

**Table 1.** Samples collected.

|  | **Breed** | **Age** |
| --- | --- | --- |
| **Dairy Cattle** | Holstein Friesian | multiparous |
| **Beef Cattle** | Simmental/Angus cross | 5-9 years old |
| **Non-lactating heifer calves** | Holstein/Angus cross | 14-16 weeks |
| **Sheep** | Polypay composite | 2-6 lactations |
| **Goat** | American alpine | 2-6 freshenings/lactations |
| **Alpaca** | Brown and Light fawn | 2-8 cria/lactations |
| **Pig** | Large White and Yorkshire cross | parity 2 or greater |
| **Human\*** | De-identified | De-identified |

\* one normal control breast tissue slide from a commercial vendor (Mopec, Madison Heights, MI, USA; Cat no. SC936) and three formalin-fixed paraffin-embedded (FFPE) archived breast tissues collected with informed consent from George Mason University provided de-identified.

### Lectin histochemistry

Mammary tissues were characterized for sialic acids using lectin histochemistry assay as previously described ^43^, for chromogenic and fluorescent staining. Briefly, formalin-fixed, paraffin-embedded tissue sections of the mammary gland were sectioned at 4 µm with placement on Superfrost® Plus slides (VWR International). Slides were dried at 60°C for 20 minutes preceding deparaffinization and staining on Roche Diagnostics’ Ventana DISCOVERY ULTRA research platform. Heat retrieval was accomplished via Cell Conditioning (CC1) solution at 100°C for 24 minutes (Roche Diagnostics Corporation). Endogenous peroxidase was quenched for chromogenic staining using Discovery Inhibitor (Roche Diagnostics Corporation). Slides were then blocked with 1X Carbo-Free blocking solution for 32 minutes (Vector Laboratories Inc.), followed by a Streptavidin/Biotin Blocking Kit (Vector Laboratories Inc.) with a separate application each of 12 minutes. The sections were incubated for 4 hours (fluorescent)/ 1 hour (chromogenic) at room temperature (RT) with one of the three lectins (SNA, MAL-I, MAL-II) from Vector Laboratories at the listed concentration in Table 2. For SA α2,6-Gal/GalNAc, slides were incubated with biotinylated SNA for both fluorescent. A cocktail of fluorescein-labeled MAL-I (specific for N-linked or O-linked glycans with SA α2,3-Galβ (1-4) GlcNAc) and biotinylated MAL-II (specific for O-linked glycans with SA α2,3-Galβ (1-3) GalNAc) were used for fluorescent staining. Biotinylated MAL-I was used separately for chromogenic staining. Following lectin incubation, streptavidin conjugated with Alexa Fluor 594 (ThermoFisher Scientific) was applied and incubated for two hours at RT. In the case of chromogenic staining, streptavidin-HRP RTU (Vector Laboratories Inc.) was incubated for 40 minutes on all three biotinylated lectins. The chromogenic signal was further amplified with DISCOVERY Amplification anti-HQ Multimer (Roche Diagnostics Corporation). Fluorescent counterstaining was performed with QD DAPI RUO (Roche Diagnostics Corporation) and mounted manually with Prolong Gold antifade mountant without DAPI (ThermoFisher Scientific). Chromogenic counterstaining was performed with hematoxylin and bluing (Roche Diagnostics Corporation), coverslipped with Sakura Tissue-Tek Film. Negative assay controls consisted of primary lectin or antibody omission. Positive lectin assay controls were performed on porcine tissues as lectin labeling has been previously established ^44^. Slides were examined and imaged using a BX-53 Olympus trinocular microscope equipped with an Olympus DP23 camera, Excelitas X-Cite mini+ compact illumination system, and CellSyns Dimension software.

**Table 2.** Lectin histochemistry and Viral binding assay reagents.

|  | Conjugate | Working Concentration | Manufacturer | Product number |
| --- | --- | --- | --- | --- |
| **Lectin Staining** |  |  |  |  |
| *Maackia amurensis* -I | Fluorescein | 20 µg/ml | Vector Laboratories | FL-1311-2 |
| *Maackia amurensis* -I | Biotinylated | 20 µg/ml | Vector Laboratories | B-1315-2 |
| *Maackia amurensis* -II | Biotinylated | 4 µg/ml | Vector Laboratories | B-1265-1 |
| *Sambucus nigra* | Biotinylated | 20 µg/ml | Vector Laboratories | B-1305-2 |
| Streptavidin | Alexa Fluor-594 | 2 µg/ml | ThermoFisher Scientific | S11227 |
| Streptavidin | Horseradish Peroxidase (HRP) - R.T.U | 1mg/ml | Vector Laboratories | SA-5704-100 |
| **Virus histochemistry** |  |  |  |  |
| Fluorescein Isothiocyanate isomer (FITC) | none | 0.1 mg/ml | Sigma-Aldrich | 3326-32-7 |
| Polyclonal Rabbit Anti-FITC antibody | HRP | 20 µg/ml | Agilent Technologies | P5100 |
| Biotin tyramide Amplification System | Streptavidin conjugated-HRP | 10 µg/ml | Akokya Biosciences | NEL700A001KT |
| 3-amino-9-ethyle-carbazol (AEC) Substrate Chromogen | none | Stock | ThermoFisher Scientific | 001122 |

### Virus preparation, inactivation, and labeling for virus histochemistry

A reverse engineered A/bald eagle/Florida/W22-134-OP/2022 (H5N1; rgA/bald eagle/FL/22) that contained an engineered low pathogenicity 2.3.4.4b H5 by removal of the polybasic cleavage site, wild type N1, and 6 internal segments from A/Puerto Rico/8/1943 (H1N1) was kindly provided by Richard Webby from St. Jude Children’s Research Hospital. Confluent layers of Madin-Darby canine kidney (MDCK) cells were inoculated with LPAIV H5N1 2.3.4.4b rgA/bald eagle/FL/22. After 48 hours, cell culture flasks went through two rounds of freeze-thaw cycles at -80°C. After the second cycle, the supernatant was harvested and cleared by low-speed centrifugation. The cleared supernatant was then centrifuged at 28,000 rpm in an SW32i rotor at 4°C for 2 hours on a layer of 10% sucrose. Pellets were resuspended using PBS buffer and measured for hemagglutination activity (HA) by 2-fold serial dilution with turkey red blood cells. The virus was then incubated at room temperature with an equal volume of 10% neutral buffered formalin for 1 hour. The inactivated virus was then dialyzed against PBS. Inactivation was confirmed on MDCK cells, and an HA was performed to determine viral titer. The inactivated virus was labeled by mixing with an equal volume of 0.1mg/ml fluorescein isothiocyanate (FITC) (Sigma-Aldrich) in 0.5 M bicarbonate buffer (pH 9.5) for 1 hour 4°C with constant agitation. The labeled virus was then dialyzed against PBS, and a final HA titer was determined.

### Viral histochemistry

Virus histochemical staining was performed on sections of lactating Holstein dairy cattle and lactating cross-bred sows. Tissues were deparaffinized with xylene and rehydrated through a series of graded alcohol solutions. A 3% hydrogen peroxide solution (Fischer Bioreagents) was used to quench endogenous peroxidases. Slides were then washed in TNT buffer (Tris (ThermoFisher Scientific), sodium chloride solution (Sigma-Aldrich), and Tween20® (Sigma-Aldrich)). Slides were then immersed in a blocking solution for 1 hour at room temperature (TNT buffer plus blocking reagent, Akoya Biosciences). Slides were incubated overnight at 4°C in a humidified chamber with FITC-labeled rgA/bald eagle/FL/22 (200 HA units per 50 µl). Tissues were then incubated in peroxidase-labeled rabbit anti-FITC antibody (Agilent Technologies) followed by signal amplification through a biotin tyramide amplification system (Akoya Biosciences). Finally, peroxidase was revealed with 3-amino-9-ethyle-carbazole (AEC, ThermoFisher Scientific), and counterstaining was performed using hematoxylin stain solution (Epredia) and ammonia hydroxide solution (Sigma-Aldrich). A sialidase control was run on sow tissue sections. Sections were pre-treated with 0.4 U/ml sialidase-A (Agilent Technologies) in 50 mM sodium phosphate buffer (pH 6.0) overnight at 37°C and subjected to the protocol as outlined above. Intestinal tissues from a chicken with and without the omission of the FITC-labeled virus were used as positive and negative controls, respectively. Slides were examined and imaged using a BX43 Olympus microscope equipped with an Olympus DP28 camera. Photomicrographs were captured using cellSens Standard.

## RESULTS

The results were broadly classified into teat and gland cistern and interlobular duct and secretory alveoli regions and summarized in Table 3.

**Table 3.** Sialic acid distribution in the mammary glands of ruminants, monogastric, and pseudo-ruminant/camelid.

| Mammary gland regions | MAL I (SA α2,3-Galβ (1-4) GlcNAc) | MAL II (SA α2,3-Galβ (1-3) GalNAc) | SNA (SA α2,6-Gal/GalNAc) |
| --- | --- | --- | --- |
| ***Ruminants (Goat/Sheep/Dairy/ Beef cattle)*** |  |  |  |
| *Teat Cistern* | + (apical; m/f) | + (apical; m/f) | + (apical; d) |
| *Gland Cistern* | + (apical; m/f) | + (apical; m/f) | + (apical; d) |
| *Interlobular (Collecting) Duct* | + (apical; * m/f) | + (apical; * m/f) | + (apical; d) |
| *Secretory Alveoli (glandular epithelium)* | +/- (apical; * m/f) | + (apical;(m/f) * to d) | + (apical; d) |
| ***Non-ruminant monogastric (Pig/Human)*** |  |  |  |
| *Teat Cistern* | + (apical; m/f)/DNE | + (apical; d)/DNE | + (apical; d)/DNE |
| *Gland Cistern* | + (apical; m/f)/DNE | + (apical; d)/DNE | + (apical; d)/DNE |
| *Interlobular (Collecting) Duct* | + (apical; m/f)/DNE | + (apical; d)/ + apical; m/f | + (apical; d)/+apical; m/f |
| *Secretory Alveoli (glandular epithelium)* | +/- (apical; m/f)/ + apical; d | + (apical; d)/+ apical; m/f | + (apical; d)/+ apical m/f |
| ***Pseudo-ruminant/camelid (Alpaca)*** |  |  |  |
| *Teat Cistern* | + (apical; m/f to d) | + (apical; m/f) | + (apical; d) |
| *Gland Cistern* | + (apical; m/f to d) | + (apical; d) | + (apical; d) |
| *Interlobular (Collecting) Duct* | + (apical; m/f to d) | + (apical; d) | + (apical; d) |
| *Secretory Alveoli (glandular epithelium)* | + (apical; m/f to d) | + (apical; d) | + (apical; d) |

Data representative of n = 3 for all animal species; n= 4 for humans. + denotes positive labeling; – no or minimal labeling; apical means labeling
lectins on the epithelial lining; m/f = multifocal labeling; d=diffuse labeling; \* denotes non-lactating cattle; DNE = did not examined.

### Lectin labeling in dairy, beef, non-lactating cattle, sheep and goat

In the ruminants (cattle, sheep, and goats), the epithelial lining of the teat (Figure 3, Panel Ai-Av) and gland cistern (Figure 3, Panel Bi-Bv) showed multifocal to diffuse, moderate positive labeling of SA α2-6 receptors (SNA) (Figure 3, Panel Ai-Av, Bi-Bv). This labeling continued into the interlobular duct (Figure 3, Ci-Cv) and secretory alveoli (Figure 3, Di-Dv) with mild to moderate intensity. There was a multifocal labeling of MAL II on the epithelial lining of the teat (Figure 3, Ei-Ev) and gland cistern (Figure 3, Fi-Fv). In the interlobular duct, MAL II labeling was rare, multifocal, and minimal to mild in intensity (Figure 3, Gi-Gv), while on the surface of secretory alveoli, the labeling was diffuse and moderate to marked in intensity (Figure 3, Hi-Hv). Interestingly, in nonlactating cattle, the intensity of MAL II labeling was comparatively low from the gland cistern (Figure 3, Fiii), and it continued to decrease in the interlobular duct and secretory alveoli (Figure 3, Giii and Hiii). Overall, the apical labeling intensity was lower for MAL II than for SNA. The spatial distribution of these receptors was apical on the epithelial cell surface and uniform across all animals examined of each species.

**Figure 3:**
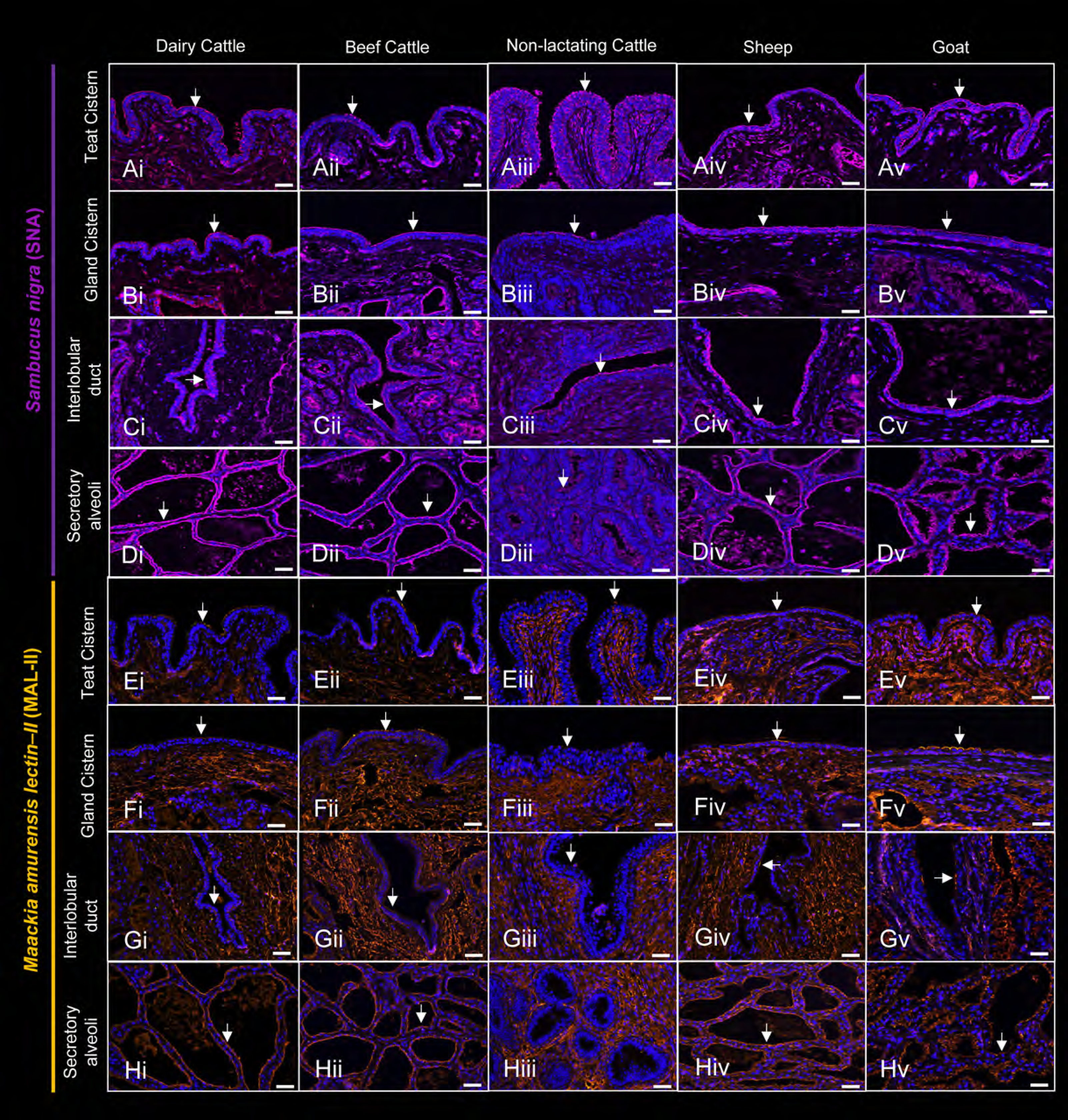
Distribution of sialic acids in the mammary gland of ruminant species. The mammary glands of dairy cattle, beef cattle, nonlactating cattle, sheep, and goats showing *Sambucus nigra lectin* (SNA, purple pseudocolor, Alexa Fluor 594) and *Maackia amurensis lectin-II* (MAL-II, red pseudocolor, Alexa Fluor 594), using fluorescent labeling. The labeling described is in reference to the epithelial surface of each image. In images Ai-Cv, there is apical, multifocal to diffuse SNA labeling in the teat cistern, gland cistern, and interlobular duct of ruminants. In Di - Dv, there is positive, diffuse, apical labeling for SNA of the epithelial surface of secretory alveoli. In images Ei-Fv, there is multifocal labeling of MAL II on the epithelial lining of the teat and gland cistern. In Gi-Gv, MAL II labeling was rare, multifocal, and minimal to mild, with the exception of the nonlactating cow in Giii, which showed minimal appreciable labeling for MAL II. In Hi-Hv, there was diffuse MAL II labeling of the epithelial surface of secretory alveoli, again with the exception of the nonlactating cow in Hiii, which also showed minimal appreciable labeling for MAL II. However, the apical labeling intensity was less for MAL II than for SNA labeling. Arrows indicating the labeling on the epithelial lining. Scale bar =50 µm.

### Lectin labeling in alpacas, pigs, and humans

Overall, the alpaca showed a similar SNA/MAL II distribution pattern to ruminants, but the labeling intensity along the epithelial lining of teat cistern, gland cistern, interlobular ducts, and secretory alveoli (Figure 4, Ai-Di and Ei-Hi), was comparatively more. This labeling on the epithelial lining is multifocal (teat cistern) to diffuse (all other tissues), intense, and apical from teat cistern to secretory alveoli. Both SNA and MAL II labeling was observed in all anatomic regions examined in the closely related monogastric species pigs and humans. In pigs, the epithelial lining of the teat cistern, gland cistern, interlobular ducts, and secretory alveoli displayed diffuse apical labeling of moderate to marked intensity for both SNA and MAL II (Figure 4, Aii – Hii). In humans, teat and gland cistern are absent hence, no images were captured (Figure 4, Aiii, Biii, Eiii, and Fiii). However, there was multifocal to diffuse apical epithelial labeling in the interlobular duct and secretory alveoli of moderate intensity for both SNA and MAL II (Figure 4, Ciii and Diii, Giii, and Hiii) in four human specimens tested.

**Figure 4:**
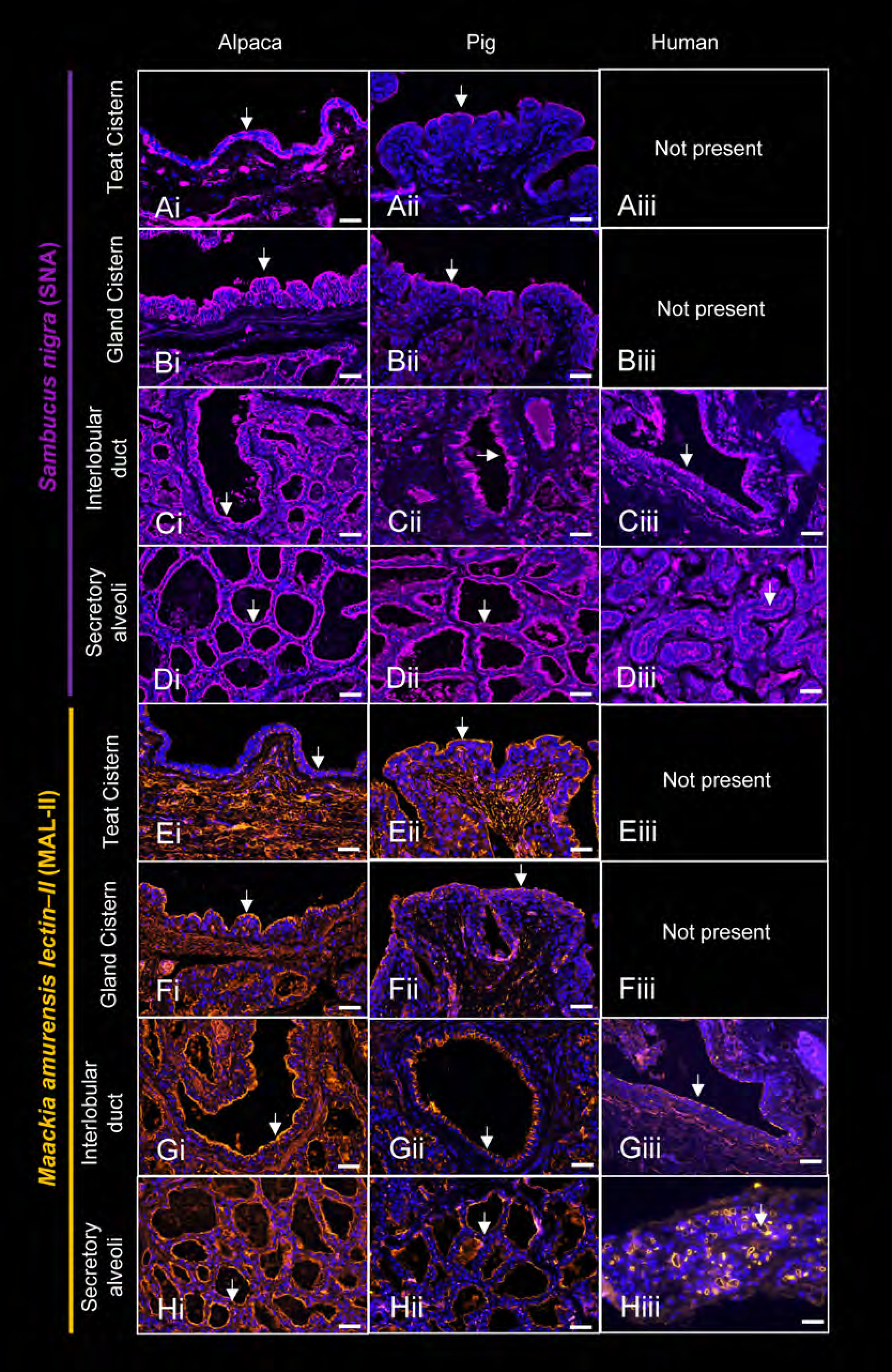
Distribution of sialic acids in the mammary gland of non-ruminant species. The mammary glands of alpaca, pig, and human showing *Sambucus nigra lectin* (SNA, purple pseudocolor, Alexa Fluor 594) and *Maackia amurensis lectin-II* (MAL-II, red pseudocolor, Alexa Fluor 594), using fluorescent labeling. In micrographs, Ai-Di, the alpaca, showed intense diffuse labeling along the epithelial lining of the teat cistern, gland cistern, interlobular duct, and secretory alveoli for SNA. Apical, multifocal, intense labeling was seen in the teat cistern for alpacas (Ei), and labeling for MAL II was diffuse, intense, and apical for MAL II on the gland cistern, interlobular ducts, and secretory alveoli (Fi, Gi, Hi). In Aii – Hii, images of pigs showed diffuse, apical labeling of moderate to marked intensity for SNA and MAL II in all anatomic regions examined. In humans, Ciii, Diii, Giii, and Hiii, a multifocal to diffuse apical epithelial labeling for both SNA and MAL II was observed in sections of the interlobular duct and secretory alveoli. Teat and gland cistern are uncommon in the human breast, hence no images, Aiii, Biii, Eiii and Fiii. Arrows indicating the labeling on the epithelial lining. Scale bar =50 µm.

### MAL I lectin labeling in ruminants and non-ruminants

The labeling of MAL I, which is specific for N-linked or O-linked glycans with SA α2,3-Galβ (1-4) GlcNAc) varied across tissue regions of mammary glands and species evaluated. In the teat and gland cistern of dairy and beef cattle, sheep, and goat, the labeling was multifocal to diffuse and moderately positive compared to occasional positive cells along the epithelial lining in non-lactating dairy cattle (Figure 5, Ai-Av and Bi -Bv). The labeling along the epithelial lining of the interlobular duct and secretory alveoli was weak positive with occasional focal labeling among all ruminants, again with non-lactating dairy cattle showing poor or no labeling (Figure 5, Ci-Cv and Di -Dv).

**Figure 5:**
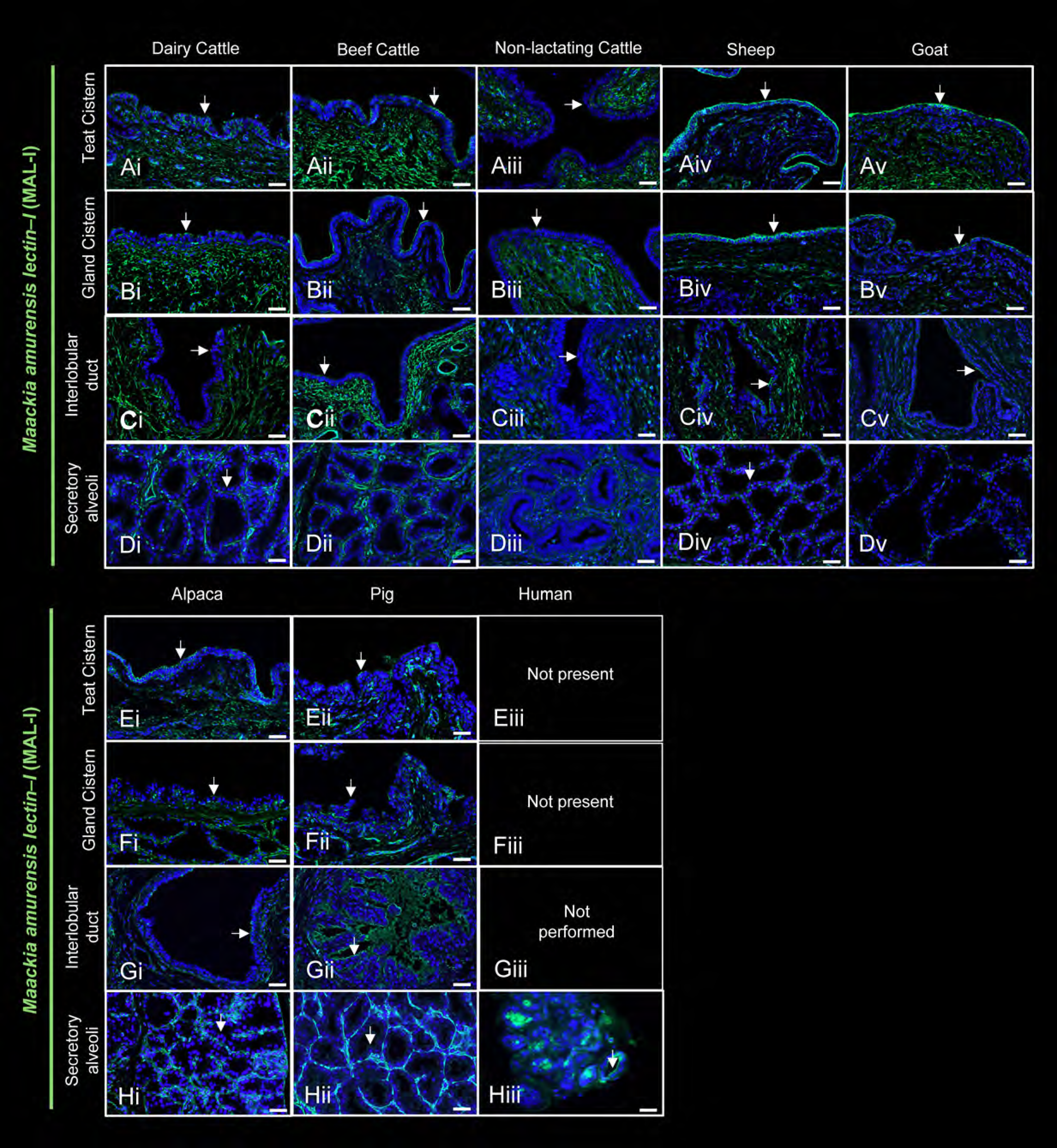
Distribution of sialic acids in the mammary gland of ruminant and non-ruminant species, MAL I specific images using IFA. The mammary glands of dairy cattle, beef cattle, nonlactating cattle, sheep, goats, alpacas, pigs, and humans showing *Maackia amurensis lectin-I* (MAL-I, green pseudocolor, Alexa Fluor 594), using fluorescent labeling. Micrographs Ai-Cv, showing apical, multifocal MAL I labeling along the epithelial lining of the teat and gland cistern, as well as the interlobular duct of ruminants. The labeling in nonlactating cattle teat and gland cistern, Aiii, and Biii, was relatively low than in other species, including the interlobular duct (Ciii). In the secretory alveoli of all ruminant species, Di - Dv, there is minimal to no apical labeling for MAL I on the epithelial surface. Ei-Gii, there is apical, multifocal MAL I labeling in the teat cistern, gland cistern, and interlobular ducts of alpacas and pigs. In the secretory alveoli of alpacas and pigs, Hi and Hii, there is apical, multifocal, minimal labeling for MAL I. There is apical, diffuse labeling for MAL I in human secretory alveoli (Hiii). Arrows indicating the labeling on the epithelial lining. Scale bar =50 µm.

In alpaca and pig, the teat and gland cistern MAL I labeling was multifocal (Figure 5, Ei-Eii and Fi-Fii). Alpacas had multifocal MAL I labeling in interlobular ducts, while pigs had continuous apical labeling (Figure 5, Gi and Gii). Like ruminants, MAL I labeling along the epithelial lining of the secretory alveoli was weak or had no labeling in alpacas and pigs (Figure 5, Hi and Hii). Meanwhile, human samples showed continuous apical labeling of MAL I in the secretory alveoli (Figure 5, Hiii). All samples were further tested using chromogenic labeling to confirm the MAL I labeling. This was done because the labeling for MAL I seemed to be less than for SNA and MAL II. Therefore, we wanted an additional assay to confirm our MAL I IFA findings. In lactating dairy cattle, multifocal labeling was seen in the teat cistern (Figure 6, Ai), with minimal to no labeling in other examined areas (Figure 6, Bi-Di). In beef cattle, there was multifocal to diffuse labeling in the teat cistern, gland cistern, and interlobular duct (Figure 6 Aii-Cii). No labeling was seen in the secretory alveoli (Figure 6m Dii). In non-lactating cattle, minimal to no labeling was noted across all anatomic locations examined (Figure 6, Aiii-Diii). For sheep and goats, there was multifocal to diffuse labeling in the teat cistern, gland cistern, and interlobular duct (Figure 6m Aiv-Civ, Av-Cv), but no labeling in the secretory alveoli in either species Figure 6 Div and Dv). In alpacas, there was multifocal to diffuse labeling for MAL I in all examined areas (Figure 6, Ei-Hi). Conversely, pigs only had MAL I labeling in the teat cistern (Figure 6, Eii). The only examined tissue for humans in this case, the interlobular duct, had diffuse apical labeling (Figure 6, Giii).

**Figure 6:**
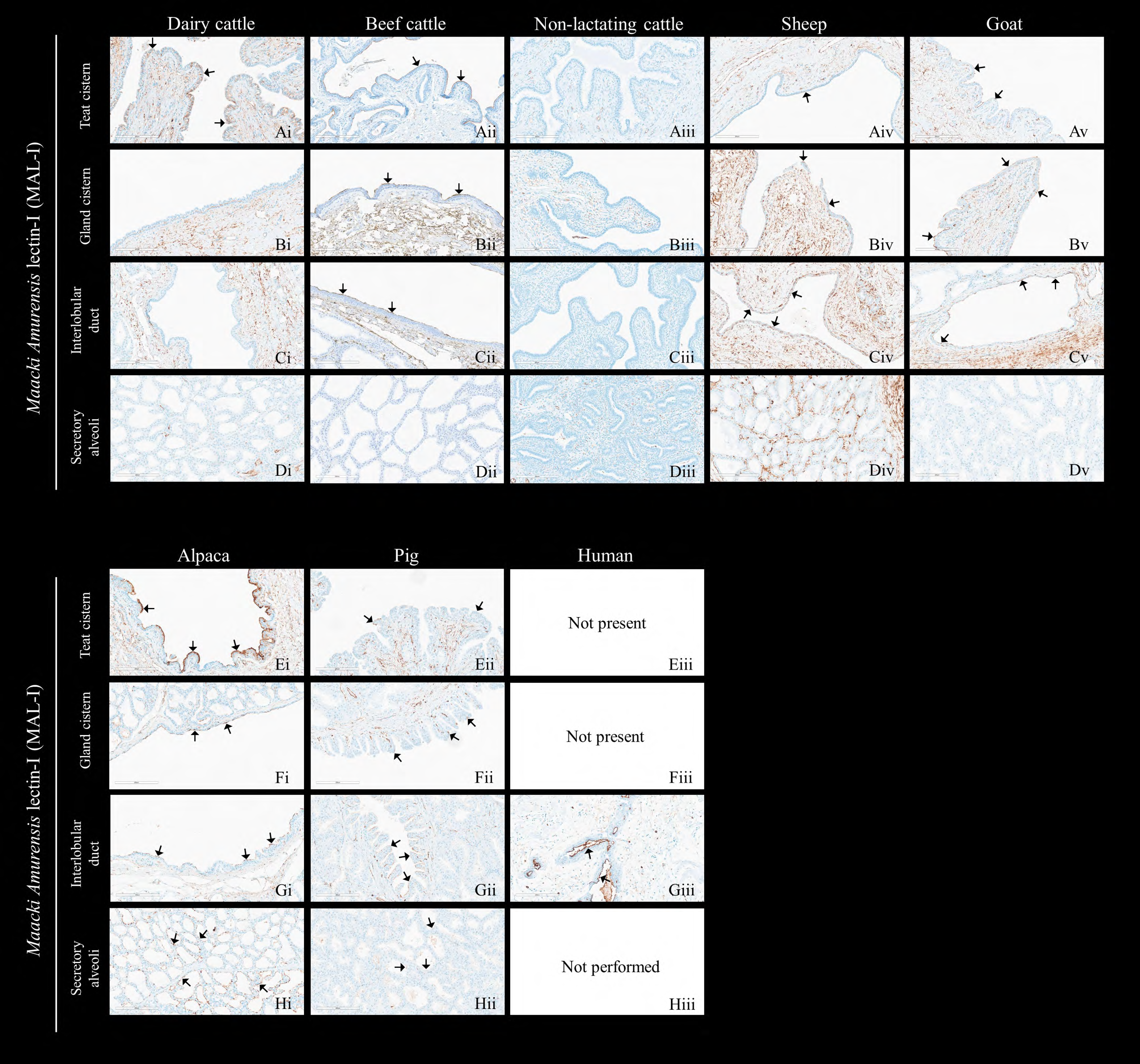
Distribution of sialic acids in the mammary gland of ruminant and non-ruminant species, MAL I specific images using chromogenic staining. The mammary glands of lactating dairy cattle, beef cattle, non-lactating cattle, sheep, and goats with *Maackia amurensis lectin-I* (MAL-I, brown chromogen), using lectin histochemistry. There is similar multifocal, apical labeling for MAL I, with no significant apical labeling observed in Aiii, Bi, Biii, Ci, and Ciii. Similarly, to the IFA technique, there is minimal labeling observed in secretory alveoli, Di-Dv. The most intense apical labeling was observed in the alpaca mammary gland, with strong apical labeling in the teat cistern, moderate labeling in the gland cistern, scant labeling in the interlobular duct, and moderate labeling within the secretory alveoli (Ei, Fi, Gi, and Hi). Faint apical labeling was observed in the teat cistern of the pig (Eii). No significant labeling was detected in the gland cistern, interlobular duct, or secretory alveoli (Fii, Gii, Hii). Strong apical labeling was present within the interlobular duct of human breast tissue (Giii). Image panel from one representative animal. Scale bar =200 µm.

### Virus histochemistry – ruminant (dairy cattle) and non-ruminant (pig)

Viral receptors, as detected by virus histochemistry, were identified at the apical epithelial surface in the interlobular ducts and secretory alveoli of both lactating Holstein cows and crossbred sows. The distribution of receptors and labeling intensity in the interlobular ducts of lactating Holstein cows varied within individual animals (Figures 7A and 7B). Labeled cells of interlobular ducts commonly had a basilar nucleus and apical cytoplasm. Cells with minimal to no labeling were commonly attenuated or not organized as a stratified cuboidal epithelial layer (Figure 7A and 7B). Intra-animal variation was also present in secretory alveoli, with some alveoli having circumferential intense labeling adjacent to secretory alveoli with no to minimal labeling to more consistent labeling of one or more epithelial cells and luminal secretions of adjacent alveoli (Figure 7C and 7D). In crossbred sows, viral receptors were commonly identified on many of the epithelial cells lining interlobular ducts, one or more epithelial cells lining secretory alveoli, and luminal secretions of adjacent alveoli (Figure 7E and 7F).

**Figure 7.**
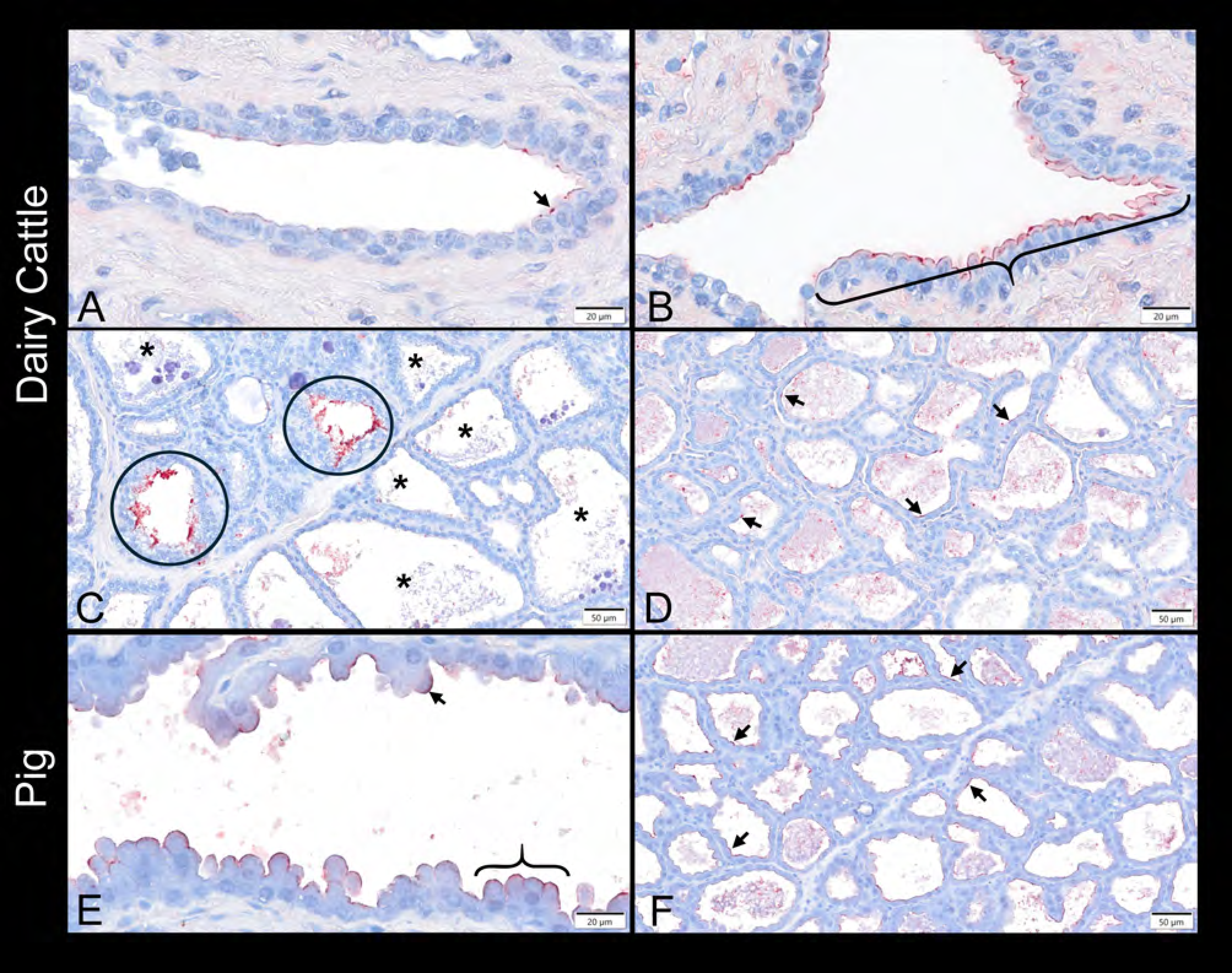
Distribution of LPAIV H5N1 2.3.4.4b rgA/bald eagle/FL/22 2.3.4.4b labeling by virus histochemistry (red) in the mammary gland of lactating Holstein cows and cross-bred sows. Interlobular ducts in the same tissue section of a Holstein dairy cow with minimal labeling (A; arrow) and intense locally extensive labeling (B; brace). Cells with minimal to no labeling were commonly attenuated (A; arrowhead) or not organized as a stratified cuboidal epithelial layer (B; arrowhead). C) Nearly circumferential intense labeling of two secretory alveoli (circled) adjacent to secretory alveoli with no to minimal labeling of the apical epithelial surface in a Holstein dairy cow (asterisk;). D) More consistent labeling of the apical surface of one or more epithelial cells (arrows) and luminal secretions of adjacent alveoli in a Holstein dairy cow. E) Interlobular duct of a crossbred sow with apical labeling of a single epithelial cell (arrow) or row of epithelial cells (brace). F) Consistent labeling of the apical surface of one or more epithelial cells (arrows) and luminal secretions of adjacent alveoli in a crossbred sow. Scale bar = 20 µm (A, B, and E); 50 µm (C, D, and F).

## DISCUSSION

As HPAI H5N1 continues to enter farms in the U.S., infecting several animals and causing human spillover events ^7,45^, this study aims to assess the risk of HPAI infecting the mammary glands of other domestic species and humans. There is a potential health risk for farm workers, and there are concerns that the virus could appear in other milk-associated products, such as goat milk or cheese, similar to the presence of HPAI H5N1 viral RNA in dairy products ^46^. In addition, many backyard farms throughout the U.S. practice mixed animal rearing, and a potential virus entry into these farms may allow for interspecies transmission, as evidenced in Oregon, USA ^47^, with the potential to mutate or reassort in these animal species. Influenza A virus infections are well known to affect the respiratory system, and it is the primary site of viral replication and subsequent pathology. But in March of 2024, the detection of HPAI H5N1 in the mammary glands of cows was indeed surprising. Identification of high levels of virus RNA in the milk and the mammary gland epithelial cells of infected dairy cattle suggested the mammary gland as a predilection site for HPAI H5N1 clade 2.3.4.4b replication in this species ^43,48^. These observations have further been confirmed through experimental studies ^10,11^. In these challenge studies, cows inoculated with HPAI H5N1 clade 2.3.4.4b viruses in the udder showed viral mastitis with epithelial necrosis within the secretory alveoli. Sequelae included exposure of the basal laminae and myoepithelial cells or replacement of epithelial cells with fibrous connective tissue ^10,11^.

One of the factors contributing to virus attachment and replication is SA, which is abundantly expressed in cow mammary gland epithelium, in particular, O-linked glycans with SA α2,3-Galβ (1-3) GalNAc, which has more affinity towards IAVs of avian origin ^43^. To the current dairy cattle outbreak, the relative abundance of both the avian and mammalian type influenza receptors on the surface of secretory epithelial cells in the mammary glands of dairy cattle most likely allowed for viral binding and entry of HPAI H5N1 2.3.4.4b into this novel anatomic area ^43^. Therefore, comparing the spatial distribution of these SA receptors in the mammary glands of ruminants and non-ruminant species, including humans, can help assess viral attachment, entry, and risk of replication in this tissue. Pathogens usually gain entry to the mammary gland through the teat canal and spread through the ducts or can be systemic and attach to the host cells with appropriate receptors. In this study, we looked at the SA along the epithelial lining of the mammary gland, starting from the teat canal to the alveoli lobules, where milk secretion occurs (Figure 1).

Although several studies have looked at the distribution of SAs in humans, pigs, cattle, sheep, goat, and camel respiratory tracts ^49,50^, to our knowledge, this is the first study to compare these receptors in the mammary glands across these species. In alpaca, testis, and placental epithelial cells, both SA α2,3-Gal and SA α2,6-Gal were present ^51^. In addition, sialic acid receptors are associated with triggering early host immune responses such as cell death signaling (necrosis/apoptosis) ^52^. Therefore, it is of utmost priority to evaluate the predisposing factors responsible for this new affinity of IAV towards mammary gland epithelium and the consequences it may have for mammalian species, including humans.

The current study showed that the mammary tissues from ruminants (cattle, sheep, and goat) and non-ruminants (alpaca, pig, and human) labeled SA α2,3-Gal and SA α2,6-Gal receptors along the mammary epithelium, suggesting that both mammalian and avian IAVs have potential to bind. The SA α2,3-Gal spatial distribution identified using MAL-I specific for N-linked or O-linked glycans with SA α2,3-Galβ (1-4) GlcNAc and MAL-II specific for O-linked glycans with SA α2,3-Galβ (1-3) GalNAc lectin labeling, was particularly interesting because of poor MAL I labeling along the epithelium (teat and gland cistern) of non-lactating cattle (Figure 5, Aiii and Biii) compared to lactating cattle. A recent study in *ex vivo* explant cultures of bovine mammary gland showed enhanced HPAI H5N1 (A/dairycattle/Texas/24-008749-001/2024) viral replication in the teat and gland cistern compared to chicken H5N1 belonging to the same 2.3.4.4b clade ^53^. Studies in mice showed selective blocking of SA α2,3-Gal associated with MAL I showed a reduction in viral replication and transmission efficiency in contact mice ^54^. Therefore, it is possible that N-linked or O-linked glycans with SA α2,3-Galβ (1-4) GlcNAc (MAL I) may play an important role in viral attachment and replication. In addition, there is also variability in lectin binding studies in the mammary gland of cattle both from the studies performed by our team and others ^43,53,55^. Further infectious and glycan biology studies are needed to confirm the role of MAL I during HPAI H5N1 viral replication in lactating dairy cattle.

This study also compared the spatial distribution of SAs in human breast tissue and showed the presence of SA α2,3-Gal and SA α2,6-Gal receptors, which agrees with recent findings ^56^. Their study further demonstrated that the proteins from HPAI H5N1 (/Texas/37/2024) bound to human mammary gland epithelium but human H1N1 and H3N2 do not bind to this tissue. The potential for HPAI H5N1 to bind to this tissue and possible replication could lead to viral mastitis, complications due to unknown etiology, breastfeeding complications, inflammation, pain, and discomfort, compromising both maternal and neonatal health. Hence, there is an immediate urgency to understand the consequences of HPAI H5N1 in human breast tissues.

The LPAIV H5N1 2.3.4.4b rgA/bald eagle/FL/22 was able to bind the apical surface of a majority of interlobular duct epithelium and a subset of epithelial cells ling secretory alveoli using virus histochemistry studies. Both structures appeared to have more consistent viral binding in the sow mammary gland than the cow mammary gland, suggesting that while not all cells are similarly susceptible to viral attachment, the sow mammary gland may contain a higher percentage of susceptible cells. The virus binding pattern partially agrees with multifocal labeling SA α2,3-Gal receptors (Table 3; Fig 3-6). However, further co-localization studies are required to determine the specificity. Pigs are susceptible to HPAI H5N1 2.3.4.4b virus strains, but whether the virus can successfully travel toward the oronasal cavity of the sow or a suckling piglet to the mammary gland is unknown ^57^. Transmission of 2009 H1N1 IAV-inoculated kits to the dam’s mammary gland has occurred ^58^. The presence of H5 receptors on the sow mammary gland suggests that attachment and replication may also be a possibility in the sow mammary gland. Endemic swine-adapted H1 and H3 IAV infection is not uncommon in suckling piglets. Yet, associated mastitis in sows has not been documented, either due to the inability of H1 or H3 IAV to cause mastitis or due to a combination of low prevalence, mild disease, and/or difficulty identifying mastitis in sows as mammary glands are not mechanically milked and not routinely evaluated without overt clinical illness. Endemic swine H1 and H3 virus histochemistry on sow mammary tissue is beyond the scope of this study.

While virus histochemistry may provide a more accurate evaluation of the presence and distribution of viral receptors than lectins or recombinant HA proteins, it has limitations. The duration of tissue fixation, tissue autolysis, and the type of fixative may alter results. In this study, livestock tissues were collected shortly after euthanasia and formalin-fixed for less than 48 hours. Additionally, the LPAIV H5N1 2.3.4.4b rgA/bald eagle/FL/22 virus was cultivated in MDCK cells. While the impact of variable virion shapes and sizes on AIV binding to receptors is not yet completely known, a virus originating from cell culture rather than a clinical specimen could impact virus binding and interpretation ^59^. In addition, the current study also showed intra-animal variability in the distribution of viral receptors based on virus histochemistry in lactating Holstein dairy cattle, suggesting that not all cells are equally susceptible to viral attachment and, therefore, infection. This variable in susceptibility is also supported by the finding that not all alveoli in inoculated animals are infected or affected by IAV and, at some level, may also contribute to the variability in the clinical disease reported from animals in the field and lesion severity of inoculated animals ^10^.

Manifestation of clinical disease is multifactorial, and while this study evaluated more Holstein dairy cattle than prior publications, it did not actively evaluate Holstein dairy cattle at different stages of lactation or known pathologic conditions that may contribute to differential glycan expression ^39,40^. The observation that epithelial cells of probable different maturation vary in labeling by virus histochemistry supports investigating differential glycan expression across different pathologic and physiological states to elucidate host factors that could influence susceptibility.

## CONCLUSION

Based on these findings, we suggest IAVs have the potential to bind to the mammary glands of cattle, sheep, goats, alpacas, pigs, and humans. Further investigation into the possible sustained IAV replication in the mammary glands of these species, both in experimental and natural environments, needs to be performed. Meanwhile, routine milk or milk product testing from these species, together with continued surveillance of the human agricultural interface, will aid in containing the virus spread, establishing mitigation strategies, and understanding evolutionary changes in IAV and their interactions with different species.

## ACKNOWLEDGMENTS

We are exceedingly grateful to Patrick Halbur, Patrick Gorden, and Rodger Main for their leadership, programmatic support, and guidance, and to the rest of the team in Iowa State University’s Veterinary Diagnostic Laboratory and Department of Veterinary Diagnostic and Production Animal Medicine. We are also grateful to Amanda Fales-Williams for her leadership and programmatic support from the Veterinary Pathology Department, in addition to the help and technical support of Amanda La Coco, Olivia Stewart, Breigha Boyle, and Katy Martin, as well as Dean Isaacson from Lab Animal Resources. Non-lactating calf mammary glands were kindly provided from archived tissues from the Matt Brewer laboratory at ISU. We also greatly appreciate the guidance, scientific advice, and support we received from Mark Ackermann of NADC, and the support of Dean Grooms, Jodi McGill, and the research support staff at the College of Veterinary Medicine at ISU.

## REFERENCES

1. Past Reported Global Human Cases with Highly Pathogenic Avian Influenza A(H5N1) (HPAI H5N1) by Country, 1997-2024 | Bird Flu | CDC [Internet]. [cited 2025 Jan 4]. Available from: https://www.cdc.gov/bird-flu/php/avian-flu-summary/chart-epi-curve-ah5n1.html

2. Shu Y, Yu H, Li D. Lethal Avian Influenza A (H5N1) Infection in a Pregnant Woman in Anhui Province, China. New England Journal of Medicine [Internet]. 2006;354(13):1421–1422. Available from: https://www.ncbi.nlm.nih.gov/pubmed/16571888 PMID: 16571888

3. Hien TT, Liem NT, Dung NT, San LT, Mai PP, Chau N van V, Suu PT, Dong VC, Mai LTQ, Thi NT, Khoa DB, Phat LP, Truong NT, Long HT, Tung CV, Giang LT, Tho ND, Nga LH, Tien NTK, San LH, Tuan L Van, Dolecek C, Thanh TT, de Jong M, Schultsz C, Cheng P, Lim W, Horby P, Farrar J. Avian Influenza A (H5N1) in 10 Patients in Vietnam. New England Journal of Medicine [Internet]. 2004/02/27. 2004;350(12):1179– 1188. Available from: https://www.ncbi.nlm.nih.gov/pubmed/14985470 PMID: 14985470

4. Van Le T, Phan LT, K Ly KH, Nguyen LT, Nguyen HT, T Ho NT, Trinh TX, Tran Minh NN, Office C, Chi Minh H, Chi Minh City Pasteur Institute H, Tuan Van Le C, Health Organization W. Fatal avian influenza A(H5N1) infection in a 36-week pregnant woman survived by her newborn in Sóc Trăng Province, Vietnam, 2012. Influenza Other Respir Viruses [Internet]. John Wiley & Sons, Ltd; 2019 May 1 [cited 2025 Feb 1];13(3):292–297. Available from: https://onlinelibrary.wiley.com/doi/full/10.1111/irv.12614 PMID: 30291769

5. WHO. Cumulative number of confirmed human cases for avian influenza A(H5N1) reported to WHO, 2003-2024, 1 November 2024. 2024.

6. Mellis AM, Coyle J, Marshall KE, Frutos AM, Singleton J, Drehoff C, Merced-Morales A, Pagano HP, Alade RO, White EB, Noble EK, Holiday C, Liu F, Jefferson S, Li ZN, Gross FL, Olsen SJ, Dugan VG, Reed C, Ellington S, Montoya S, Kohnen A, Stringer G, Alden N, Blank P, Chia D, Bagdasarian N, Herlihy R, Lyon-Callo S, Levine MZ. Serologic Evidence of Recent Infection with Highly Pathogenic Avian Influenza A(H5) Virus Among Dairy Workers — Michigan and Colorado, June– August 2024. MMWR Morb Mortal Wkly Rep [Internet]. 2024 Nov 7 [cited 2025 Jan 4];73(44):1004–1009. Available from: https://www.cdc.gov/mmwr/volumes/73/wr/mm7344a3.htm

7. CDC. H5 Bird Flu: Current Situation | Bird Flu | CDC [Internet]. 2025 [cited 2025 Jan 27]. Available from: https://www.cdc.gov/bird-flu/situation-summary/index.html

8. Uiprasertkul M, Puthavathana P, Sangsiriwut K, Pooruk P, Srisook K, Peiris M, Nicholls JM, Chokephaibulkit K, Vanprapar N, Auewarakul P. Influenza A H5N1 replication sites in humans. Emerg Infect Dis [Internet]. 2005;11(7):1036–1041. Available from: https://www.ncbi.nlm.nih.gov/pubmed/16022777 PMID: 16022777

9. Naguib MM, Eriksson P, Jax E, Wille M, Lindskog C, Bröjer C, Krambrich J, Waldenström J, Kraus RHS, Larson G, Lundkvist Å, Olsen B, Järhult JD, Ellström P. A Comparison of Host Responses to Infection with Wild-Type Avian Influenza Viruses in Chickens and Tufted Ducks. Microbiol Spectr [Internet]. Microbiol Spectr; 2023 Aug 17 [cited 2025 Feb 27];11(4). Available from: https://pubmed.ncbi.nlm.nih.gov/37358408/ PMID: 37358408

10. Baker AL, Arruda B, Palmer M V., Boggiatto P, Sarlo Davila K, Buckley A, Ciacci Zanella G, Snyder CA, Anderson TK, Hutter CR, Nguyen TQ, Markin A, Lantz K, Posey EA, Kim Torchetti M, Robbe-Austerman S, Magstadt DR, Gorden PJ. Dairy cows inoculated with highly pathogenic avian influenza virus H5N1. Nature [Internet]. Nature Publishing Group; 2024 Oct 15 [cited 2024 Nov 6];1–3. Available from: https://www.nature.com/articles/s41586-024-08166-6 PMID: 39406346

11. Halwe NJ, Cool K, Breithaupt A, Schön J, Trujillo JD, Nooruzzaman M, Kwon T, Ahrens AK, Britzke T, McDowell CD, Piesche R, Singh G, Pinho dos Reis V, Kafle S, Pohlmann A, Gaudreault NN, Corleis B, Ferreyra FM, Carossino M, Balasuriya UBR, Hensley L, Morozov I, Covaleda LM, Diel D, Ulrich L, Hoffmann D, Beer M, Richt JA. H5N1 clade 2.3.4.4b dynamics in experimentally infected calves and cows. Nature 2024 [Internet]. Nature Publishing Group; 2024 Sep 25 [cited 2024 Oct 14];1–3. Available from: https://www.nature.com/articles/s41586-024-08063-y

12. USDA-APHIS-Wildlife Services. HPAI Detections in Mammals [Internet]. National Wildlife Disease Program. 2025 [cited 2025 Mar 13]. Available from: https://www.aphis.usda.gov/livestock-poultry-disease/avian/avian-influenza/hpai-detections/mammals

13. Schlaudecker EP, Steinhoff MC, Omer SB, McNeal MM, Roy E, Arifeen SE, Dodd CN, Raqib R, Breiman RF, Zaman K. IgA and neutralizing antibodies to influenza a virus in human milk: a randomized trial of antenatal influenza immunization. PLoS One [Internet]. PLoS One; 2013 Aug 14 [cited 2025 Feb 27];8(8). Available from: https://pubmed.ncbi.nlm.nih.gov/23967126/ PMID: 23967126

14. Bigler NA, Bruckmaier RM, Gross JJ. Implications of placentation type on species-specific colostrum properties in mammals. J Anim Sci [Internet]. J Anim Sci; 2022 Dec 1 [cited 2025 Feb 27];100(12). Available from: https://pubmed.ncbi.nlm.nih.gov/36048628/ PMID: 36048628

15. Walther B, Guggisberg D, Badertscher R, Egger L, Portmann R, Dubois S, Haldimann M, Kopf-Bolanz K, Rhyn P, Zoller O, Veraguth R, Rezzi S. Comparison of nutritional composition between plant-based drinks and cow’s milk. Front Nutr [Internet]. Front Nutr; 2022 Oct 28 [cited 2025 Feb 27];9. Available from: https://pubmed.ncbi.nlm.nih.gov/36386959/ PMID: 36386959

16. Macias H, Hinck L. Mammary gland development. Wiley Interdiscip Rev Dev Biol [Internet]. Wiley Interdiscip Rev Dev Biol; 2012 Jul [cited 2025 Feb 27];1(4):533–557. Available from: https://pubmed.ncbi.nlm.nih.gov/22844349/ PMID: 22844349

17. Cimmino F, Catapano A, Villano I, Di Maio G, Petrella L, Traina G, Pizzella A, Tudisco R, Cavaliere G. Invited review: Human, cow, and donkey milk comparison: Focus on metabolic effects. J Dairy Sci. Elsevier; 2023 May 1;106(5):3072–3085. PMID: 36894420

18. Haxhiaj K, Wishart DS, Ametaj BN. Mastitis: What It Is, Current Diagnostics, and the Potential of Metabolomics to Identify New Predictive Biomarkers. Dairy 2022, Vol 3, Pages 722-746 [Internet]. Multidisciplinary Digital Publishing Institute; 2022 Oct 17 [cited 2025 Feb 27];3(4):722–746. Available from: https://www.mdpi.com/2624-862X/3/4/50/htm

19. Schukken YH, Günther J, Fitzpatrick J, Fontaine MC, Goetze L, Holst O, Leigh J, Petzl W, Schuberth HJ, Sipka A, Smith DGE, Quesnell R, Watts J, Yancey R, Zerbe H, Gurjar A, Zadoks RN, Seyfert HM. Host-response patterns of intramammary infections in dairy cows. Vet Immunol Immunopathol. Elsevier; 2011 Dec 15;144(3–4):270–289. PMID: 21955443

20. Mastitis in Cattle - Reproductive System - Merck Veterinary Manual [Internet]. [cited 2025 Feb 27]. Available from: https://www.merckvetmanual.com/reproductive-system/mastitis-in-large-animals/mastitis-in-cattle

21. Matrosovich M, Zhou N, Kawaoka Y, Webster R. The Surface Glycoproteins of H5 Influenza Viruses Isolated from Humans, Chickens, and Wild Aquatic Birds Have Distinguishable Properties. J Virol [Internet]. American Society for Microbiology; 1999 Feb [cited 2024 Dec 2];73(2):1146–1155. Available from: https://journals.asm.org/doi/10.1128/jvi.73.2.1146-1155.1999 PMID: 9882316

22. Matrosovich M, Herrler G, Klenk HD. Sialic Acid Receptors of Viruses. In: Gerardy-Schahn R, Delannoy P, Von Itzstein M, editors. SialoGlyco Chemistry and Biology II [Internet]. Cham: Springer International Publishing; 2013. p. 1–28. Available from: http://link.springer.com/10.1007/128_2013_466

23. Nguyen L, McCord KA, Bui DT, Bouwman KM, Kitova EN, Elaish M, Kumawat D, Daskhan GC, Tomris I, Han L, Chopra P, Yang TJ, Willows SD, Mason AL, Mahal LK, Lowary TL, West LJ, Hsu STD, Hobman T, Tompkins SM, Boons GJ, de Vries RP, Macauley MS, Klassen JS. Sialic acid-containing glycolipids mediate binding and viral entry of SARS-CoV-2. Nat Chem Biol [Internet]. Nature Publishing Group; 2022 Nov 9 [cited 2024 Oct 17];18(1):81–90. Available from: https://www.nature.com/articles/s41589-021-00924-1 PMID: 34754101

24. Tan CW, Huan Hor CH, Kwek S Sen, Tee HK, Sam IC, Goh ELK, Ooi EE, Chan YF, Wang LF. Cell surface α2,3-linked sialic acid facilitates Zika virus internalization. Emerg Microbes Infect [Internet]. Taylor and Francis Ltd.; 2019 Jan 1 [cited 2024 Oct 17];8(1):426–437. Available from: https://pmc.ncbi.nlm.nih.gov/articles/PMC6455136/ PMID: 30898036

25. Taube S, Perry JW, Yetming K, Patel SP, Auble H, Shu L, Nawar HF, Lee CH, Connell TD, Shayman JA, Wobus CE. Ganglioside-Linked Terminal Sialic Acid Moieties on Murine Macrophages Function as Attachment Receptors for Murine Noroviruses. J Virol [Internet]. American Society for Microbiology; 2009 May [cited 2024 Oct 17];83(9):4092–4101. Available from: https://pmc.ncbi.nlm.nih.gov/articles/PMC2668497/ PMID: 19244326

26. Haselhorst T, Fleming FE, Dyason JC, Hartnell RD, Yu X, Holloway G, Santegoets K, Kiefel MJ, Blanchard H, Coulson BS, Von Itzstein M. Sialic acid dependence in rotavirus host cell invasion. Nat Chem Biol [Internet]. Nature Publishing Group; 2009 Dec 21 [cited 2024 Oct 17];5(2):91–93. Available from: https://www.nature.com/articles/nchembio.134 PMID: 19109595

27. Huang LY, Patel A, Ng R, Miller EB, Halder S, McKenna R, Asokan A, Agbandje-McKenna M. Characterization of the Adeno-Associated Virus 1 and 6 Sialic Acid Binding Site. J Virol [Internet]. American Society for Microbiology; 2016 Jun [cited 2024 Oct 17];90(11):5219. Available from: https://pmc.ncbi.nlm.nih.gov/articles/PMC4934739/ PMID: 26962225

28. Arnberg N, Edlund K, Kidd AH, Wadell G. Adenovirus Type 37 Uses Sialic Acid as a Cellular Receptor. J Virol [Internet]. American Society for Microbiology; 2000 Jan [cited 2024 Oct 17];74(1):42–48. Available from: https://pmc.ncbi.nlm.nih.gov/articles/PMC111511/ PMID: 10590089

29. Zhang Y, Liu W, He F, Liu YJ, Jiang H, Hao C, Wang W. Myosin 9 and N-glycans jointly regulate human papillomavirus entry. Journal of Biological Chemistry [Internet]. American Society for Biochemistry and Molecular Biology Inc.; 2024 Feb 1 [cited 2024 Oct 17];300(2). Available from: http://www.jbc.org/article/S002192582400036X/fulltext PMID: 38242322

30. O’Hara SD, Stehle T, Garcea R. Glycan receptors of the Polyomaviridae: Structure, function, and pathogenesis. Curr Opin Virol. Elsevier; 2014 Aug 1;7(1):73–78. PMID: 24983512

31. Stuart AD, Brown TDK. α2,6-linked sialic acid acts as a receptor for Feline calicivirus. Journal of General Virology [Internet]. Microbiology Society; 2007 Jan 1 [cited 2024 Oct 17];88(1):177–186. Available from: https://www.microbiologyresearch.org/content/journal/jgv/10.1099/vir.0.82158-0 PMID: 17170450

32. Hopkins AP, Hawkhead JA, Thomas GH. Transport and catabolism of the sialic acids N-glycolylneuraminic acid and 3-keto-3-deoxy-d-glycero-d-galactonononic acid by Escherichia coli K-12. FEMS Microbiol Lett [Internet]. Oxford Academic; 2013 Oct 1 [cited 2024 Oct 17];347(1):14–22. Available from: 10.1111/1574-6968.12213 PMID: 23848303

33. Rifatbegović M, Nicholas RAJ, Mutevelić T, Hadžiomerović M, Maksimović Z. Pathogens Associated with Bovine Mastitis: The Experience of Bosnia and Herzegovina. Vet Sci [Internet]. Multidisciplinary Digital Publishing Institute (MDPI); 2024 Feb 1 [cited 2024 Oct 17];11(2):63. Available from: https://pmc.ncbi.nlm.nih.gov/articles/PMC10891550/

34. Eneva R, Engibarov S, Abrashev R, Krumova E, Angelova M. Sialic acids, sialoconjugates and enzymes of their metabolism in fungi. Biotechnology and Biotechnological Equipment [Internet]. Taylor & Francis; 2021 [cited 2024 Oct 17];35(1):364–375. Available from: https://www.tandfonline.com/doi/abs/10.1080/13102818.2021.1879678

35. Cavalcante T, Medeiros MM, Mule SN, Palmisano G, Stolf BS. The Role of Sialic Acids in the Establishment of Infections by Pathogens, With Special Focus on Leishmania. Front Cell Infect Microbiol [Internet]. Frontiers Media S.A.; 2021 May 13 [cited 2024 Oct 17];11:671913. Available from: www.frontiersin.org PMID: 34055669

36. Warwas ML, Watson JN, Bennet AJ, Moore MM. Structure and role of sialic acids on the surface of Aspergillus fumigatus conidiospores. Glycobiology [Internet]. Oxford Academic; 2007 Apr 1 [cited 2024 Oct 17];17(4):401–410. Available from: 10.1093/glycob/cwl085 PMID: 17223648

37. Sharma R, Ahlawat S, Sharma H, Aggarwal RAK, Sharma V, Tantia MS. Variable sialic acid content in milk of Indian cattle and buffalo across different stages of lactation. Journal of Dairy Research [Internet]. 2019 Feb;86(1):98–101. Available from: https://www.cambridge.org/core/product/identifier/S002202991800081X/type/journal_article

38. Wang B, Brand-Miller J. The role and potential of sialic acid in human nutrition. European Journal of Clinical Nutrition 2003 57:11 [Internet]. Nature Publishing Group; 2003 Oct 23 [cited 2025 Feb 27];57(11):1351–1369. Available from: https://www.nature.com/articles/1601704 PMID: 14576748

39. de Sousa YRF, da Silva Vasconcelos MA, Costa RG, de Azevedo Filho CA, de Paiva EP, Queiroga R de CR do E. Sialic acid content of goat milk during lactation. Livest Sci [Internet]. 2015 Jul;177:175–180. Available from: https://linkinghub.elsevier.com/retrieve/pii/S1871141315001869

40. Puentea R, Huesob P. Lactational Changes in the N-Glycoloylneuraminic Acid Content of Bovine Milk Gangliosides. Biol Chem Hoppe Seyler [Internet]. 1993 Jan;374(7–12):475–478. Available from: https://www.degruyter.com/document/doi/10.1515/bchm3.1993.374.7-12.475/html PMID: 8216898

41. Zhao C, Hu X, Qiu M, Bao L, Wu K, Meng X, Zhao Y, Feng L, Duan S, He Y, Zhang N, Fu Y. Sialic acid exacerbates gut dysbiosis-associated mastitis through the microbiota-gut-mammary axis by fueling gut microbiota disruption. Microbiome [Internet]. BioMed Central Ltd; 2023 Dec 1 [cited 2024 Oct 17];11(1):78. Available from: https://pmc.ncbi.nlm.nih.gov/articles/PMC10107595/ PMID: 37069691

42. Teoh ST, Ogrodzinski MP, Ross C, Hunter KW, Lunt SY. Sialic acid metabolism: A key player in breast cancer metastasis revealed by metabolomics. Front Oncol [Internet]. Frontiers Media S.A.; 2018 May 28 [cited 2025 Jan 4];8(MAY):360894. Available from: www.frontiersin.org

43. Nelli RK, Harm TA, Siepker C, Groeltz-Thrush JM, Jones B, Twu NC, Nenninger AS, Magstadt DR, Burrough ER, Piñeyro PE, Mainenti M, Carnaccini S, Plummer PJ, Bell TM. Sialic Acid Receptor Specificity in Mammary Gland of Dairy Cattle Infected with Highly Pathogenic Avian Influenza A(H5N1) Virus. Emerg Infect Dis. Centers for Disease Control and Prevention (CDC); 2024 Jul 1;30(7):1361–1373. PMID: 38861554

44. Nelli RK, Kuchipudi S V, White GA, Perez BB, Dunham SP, Chang KC. Comparative distribution of human and avian type sialic acid influenza receptors in the pig. BMC Vet Res [Internet]. 2010 Dec;6(1):4. Available from: https://bmcvetres.biomedcentral.com/articles/10.1186/1746-6148-6-4

45. 2022–2024 Detections of Highly Pathogenic Avian Influenza [Internet]. [cited 2025 Jan 27]. Available from: https://www.aphis.usda.gov/livestock-poultry-disease/avian/avian-influenza/hpai-detections

46. Suarez DL, Goraichuk I V., Killmaster L, Spackman E, Clausen NJ, Colonius TJ, Leonard CL, Metz ML. Testing of Retail Cheese, Butter, Ice Cream, and Other Dairy Products for Highly Pathogenic Avian Influenza in the US. J Food Prot [Internet]. Elsevier; 2025 Jan 2 [cited 2025 Jan 27];88(1):100431. Available from: https://linkinghub.elsevier.com/retrieve/pii/S0362028X24002151

47. USDA Animal and Plant Health Inspection Service Shares Update on H5N1 Detection in Oregon Swine, Bovine Vaccine Candidate Progression | Animal and Plant Health Inspection Service [Internet]. [cited 2025 Jan 27]. Available from: https://www.aphis.usda.gov/news/agency-announcements/usda-animal-plant-health-inspection-service-shares-update-h5n1-detection

48. Burrough ER, Magstadt DR, Petersen B, Timmermans SJ, Gauger PC, Zhang J, Siepker C, Mainenti M, Li G, Thompson AC, Gorden PJ, Plummer PJ, Main R. Highly Pathogenic Avian Influenza A(H5N1) Clade 2.3.4.4b Virus Infection in Domestic Dairy Cattle and Cats, United States, 2024. Emerg Infect Dis [Internet]. Centers for Disease Control and Prevention (CDC); 2024 Jul 1 [cited 2024 Jul 22];30(7):1335–1343. Available from: https://wwwnc.cdc.gov/eid/article/30/7/24-0508_article PMID: 38683888

49. Li W, Hulswit RJG, Widjaja I, Raj VS, McBride R, Peng W, Widagdo W, Tortorici MA, Van Dieren B, Lang Y, Van Lent JWM, Paulson JC, De Haan CAM, De Groot RJ, Van Kuppeveld FJM, Haagmans BL, Bosch BJ. Identification of sialic acid-binding function for the Middle East respiratory syndrome coronavirus spike glycoprotein. Proc Natl Acad Sci U S A [Internet]. National Academy of Sciences; 2017 Oct 3 [cited 2025 Jan 4];114(40):E8508–E8517. Available from: https://www.pnas.org/doi/abs/10.1073/pnas.1712592114 PMID: 28923942

50. Kuchipudi S V., Nelli RK, Gontu A, Satyakumar R, Nair MS, Subbiah M. Sialic acid receptors: The key to solving the enigma of zoonotic virus spillover. Viruses [Internet]. Multidisciplinary Digital Publishing Institute; 2021 Feb 8 [cited 2021 Feb 8];13(2):262. Available from: 10.3390/v13020262 PMID: 33567791

51. Parillo F, Magi GE, Diverio S, Catone G. Immunohistochemical and lectin histochemical analysis of the alpaca efferent ducts. Histol Histopathol [Internet]. Histol Histopathol; 2009 [cited 2024 Nov 11];24(1):1–12. Available from: https://pubmed.ncbi.nlm.nih.gov/19012239/ PMID: 19012239

52. Malagolini N, Chiricolo M, Marini M, Dall’Olio F. Exposure of α2,6-sialylated lactosaminic chains marks apoptotic and necrotic death in different cell types. Glycobiology [Internet]. Oxford Academic; 2009 Feb 1 [cited 2024 Nov 10];19(2):172–181. Available from: 10.1093/glycob/cwn122 PMID: 18988689

53. Imai M, Ueki H, Ito M, Iwatsuki-Horimoto K, Kiso M, Biswas A, Trifkovic S, Cook N, Halfmann PJ, Neumann G, Eisfeld AJ, Kawaoka Y. Highly pathogenic avian H5N1 influenza A virus replication in ex vivo cultures of bovine mammary gland and teat tissues. Emerg Microbes Infect [Internet]. Taylor and Francis Ltd.; 2025 [cited 2025 Mar 3];14(1). Available from: https://www.tandfonline.com/action/journalInformation?journalCode=temi20 PMID: 39781889

54. Ortigoza MB, Mobini CL, Rocha HL, Bartlett S, Loomis CA, Weiser JN. Inhibiting influenza virus transmission using a broadly acting neuraminidase that targets host sialic acids in the upper respiratory tract. mBio [Internet]. American Society for Microbiology; 2024 Feb 1 [cited 2025 Mar 3];15(2). Available from: https://journals.asm.org/doi/10.1128/mbio.02203-23 PMID: 38206008

55. Kristensen C, Larsen LE, Trebbien R, Jensen HE. The avian influenza A virus receptor SA-α2,3-Gal is expressed in the porcine nasal mucosa sustaining the pig as a mixing vessel for new influenza viruses. Virus Res. Elsevier B.V.; 2024 Feb 1;340. PMID: 38142890

56. Song H, Hao T, Han P, Wang H, Zhang X, Li X, Wang Y, Chen J, Li Y, Jin X, Duan X, Zhang W, Bi Y, Jin R, Sun L, Wang N, Gao GF. Receptor binding, structure, and tissue tropism of cattle-infecting H5N1 avian influenza virus hemagglutinin. Cell [Internet]. Elsevier B.V.; 2025 Feb 20 [cited 2025 Mar 3];188(4):919–929.e9. Available from: https://www.cell.com/action/showFullText?pii=S0092867425000480

57. Arruda B, Vincent Baker AL, Buckley A, Anderson TK, Torchetti M, Bergeson NH, Killian ML, Lantz K. Divergent Pathogenesis and Transmission of Highly Pathogenic Avian Influenza A(H5N1) in Swine - Volume 30, Number 4—April 2024 - Emerging Infectious Diseases journal - CDC. Emerg Infect Dis [Internet]. Centers for Disease Control and Prevention (CDC); 2024 Apr 1 [cited 2025 Mar 3];30(4):738–751. Available from: https://wwwnc.cdc.gov/eid/article/30/4/23-1141_article PMID: 38478379

58. Paquette SG, Banner D, Huang SSH, Almansa R, Leon A, Xu L, Bartoszko J, Kelvin DJ, Kelvin AA. Influenza Transmission in the Mother-Infant Dyad Leads to Severe Disease, Mammary Gland Infection, and Pathogenesis by Regulating Host Responses. PLoS Pathog [Internet]. Public Library of Science; 2015 [cited 2025 Mar 3];11(10):e1005173. Available from: https://journals.plos.org/plospathogens/article?id=10.1371/journal.ppat.1005173 PMID: 26448646

59. Partlow EA, Jaeggi-Wong A, Planitzer SD, Berg N, Li Z, Ivanovic T. Influenza A virus rapidly adapts particle shape to environmental pressures. Nature Microbiology 2025 10:3 [Internet]. Nature Publishing Group; 2025 Feb 10 [cited 2025 Mar 3];10(3):784–794. Available from: https://www.nature.com/articles/s41564-025-01925-9

